# Globally Harmonized Carbon Storage Data

**DOI:** 10.1101/727750

**Authors:** Justin Andrew Johnson

## Abstract

This paper presents methods and results that combine multiple carbon storage and above-ground biomass datasets using a simple decision tree approach. The resulting dataset combines the positive attributes of the multiple input datasets in order to have global, high-resolution extent while utilizing the best statistical methods where possible. Visual inspection shows very different spatial configurations of carbon storage between the results and the input data, suggesting that combination of methods can improve estimates. The summation of the decision tree result was 336.55 petagrams while the summation of a dataset based on the IPCC Tier 1 method was 502.38 petagrams (49.27% higher than the results from the decision tree).

## Introduction

Many analyses of global sustainability assess how climate change will be affected by carbon storage and sequestration. The primary method for estimating how carbon will be stored on a landscape, the IPCC Tier 1 method, assigns a single value for carbon storage to entire land-use, land-cover classes across large ecoregions. Recent advances based on remote sensing show that there is considerable variation in carbon storage, even within a localized area (discussed in depth below). However, these studies have had a number of shortcomings, namely being focused only on pantropic regions or only on forest-cover. Additionally, there exists differences among these new approaches, both in terms of statistical methods and in terms of the resulting carbon storage values attributed to the landscape. Finally, basic tasks of reprojection, resampling and converting to similar units (carbon content instead of above-ground biomass) need to be done in a consistent way for valid comparisons among datasets. This paper presents a simple decision tree approach to combine the different datasets in a way that provides globally harmonized values using the best available data for all locations. The final result is a global-extent map of carbon storage per hectare, reported at 30 arcsecond resolution (slightly less than 1km at the equator).

### Existing estimates of carbon storage

The first broadly used, global extent dataset on carbon storage was produced by Ruesch and Gibbs (2008). Based on default values from the IPCC Good Practice Guidance (Penman et al. 2003) and the Greenhosue Gas Inventory Guidelines (IPCC 2006), Ruesch and Gibbs synthesized and mapped the values to the Global Land Cover 2000 Project (GLC2000) land-use, land-cover (LULC) map. This approach was described as a “paint by numbers” approach insofar as it assigned a single carbon value to all locations based on their LULC class and ecoregion. These data are represented in figure 1, column A.

**Figure 1:**
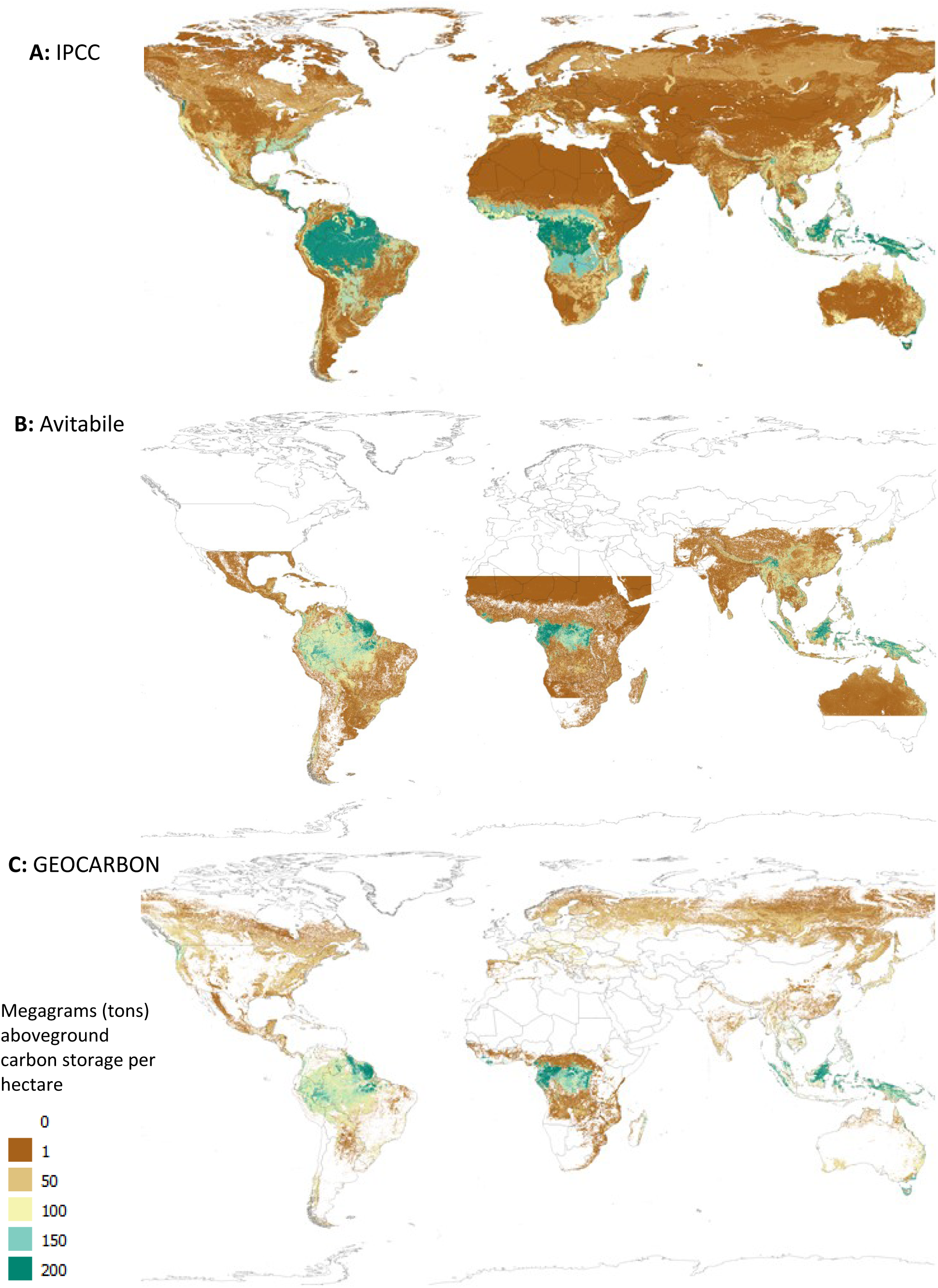
Aboveground carbon storage from three existing datasets.

New estimates of carbon storage have emerged that are based on statistically linking remote sensing data with field observations of carbon storage. One of the first of these, Saatchi et al. (2011), combined 4079 field observations with LiDAR, optical and microwave imagery in a maximum entropy regression. This generated extrapolations over the landscape at 1 km resolution. Shortly after the release of the Saatchi data, Baccini et al. (2012) released a dataset with similar characteristics but different input data and methods. Baccini et al used LiDAR, field data and allometric relationships, couple with MODIS imagery to produce 500 m data. These two datasets both exhibited very large improvements in accuracy over the Ruesch and Gibbs method, but they differed from each other considerably.

Avitabile et al. (2016) addressed the discrepancies between these data using a statistical fusion method, along with additional input data, to produce a product that improved upon both the Saatchi and Baccini data (see figure 1B). Avitabile et al. compiled ground observations and locally calibrated high-resolution biomass maps to create a reference dataset which was then used to optimally combine the Saatchi and Baccini data using data-fusion methods from Ge et al. (2014). Compared to both input maps, the Avitabile product improved accuracy in a number of locations, showing higher aboveground biomass density in the Congo basin, Eastern Amazon and South-East Asia, and lower values in much of Central America and dryland areas of Africa. The validation exercise showed a root mean squared error (RMSE) 15-21% lower than the input maps. Additionally, the estimates, when compared to the validation field plots, were no longer biased on average. For these reasons, I use Avitabile as a key input to the decision tree method presented here and do not directly incorporate Saatchi and Baccini. Note, however, that Avitabile was 1km resolution (slightly more than 30 arcseconds) and pan-tropical.

Several analyses consider carbon storage values outside of the tropics, though this area remains relatively less-studied. Santoro et al. (2015) presented a was outside the tropics (north of 10 degrees) that used hyper-temporal Envisat ASAR using the BIOMASAR algorithm to report aboveground biomass estimates at 0.01 arcdegrees. However, these calculations were only valid for forest areas, thus reporting zero values for any area not in a GLC2000 forest class. Shortly upon release, the GEOCARBON (Avitabile et al. 2014) project further synthesized this product, combining it with the Avitabile data, thereby providing a global map of carbon storage, though only for forest LULC classes. This is the second key input in the decision tree and is presented in figure 1C.

## Methods

Of the datasets described above, only the Ruesch and Gibbs dataset has both global coverage and includes all LULC classes, though this dataset lacks calibration to remote sensing and field data. This section describes how I combined the different datasets via a simple decision tree. Several of the datasets have already been combined, namely via the data-fusion of Avitabile and the GEOCARBON work to incorporate non-tropical forests. Using these inputs, the decision rule is simply:

1. If in the tropics and Avitabile is non-zero, use it.
2. In remaining places, where Santoro/GEOCARBON is zero, use it.
3. In remaining places, use Ruesch and Gibbs values.

Prior to applying the decision tree, all maps were reprojected to 30 arcseconds. Note that this is different than all of the inputs, but is very close to the Avitabile resolution (Avitabile used 1 km, or approximately 32.04 arcseconds. The 30 arcsecond resolution was chosen because it is close to Avitabile but is an even fraction of arcdegrees (and thus can more easily be aggregated to other scales).

Also prior to the final calculation, all datasets were converted to carbon storage values rather than above-ground biomass (AGB) estimates. There exist competing assumption for how best to do this. For example, Saatchi et al. used the a conversion coefficient of 0.5 to convert from AGB to carbon. However, for this approach, I followed the methods presented in Djomo et al. (2011) as summarized by Vashum and Jayakumar (2012). Djomo et al. analyzed the carbon content of wood in a CNS analyzer and found a mean value of 46.53%. Based on this, I converted all AGB to carbon storage by multiplying by 0.4653.

## Results

Applying the data conversions and decision tree described above generated results that were global in extent and was not missing non-forest LULC classes. The results are shown in figure 2.

**Figure 2:**
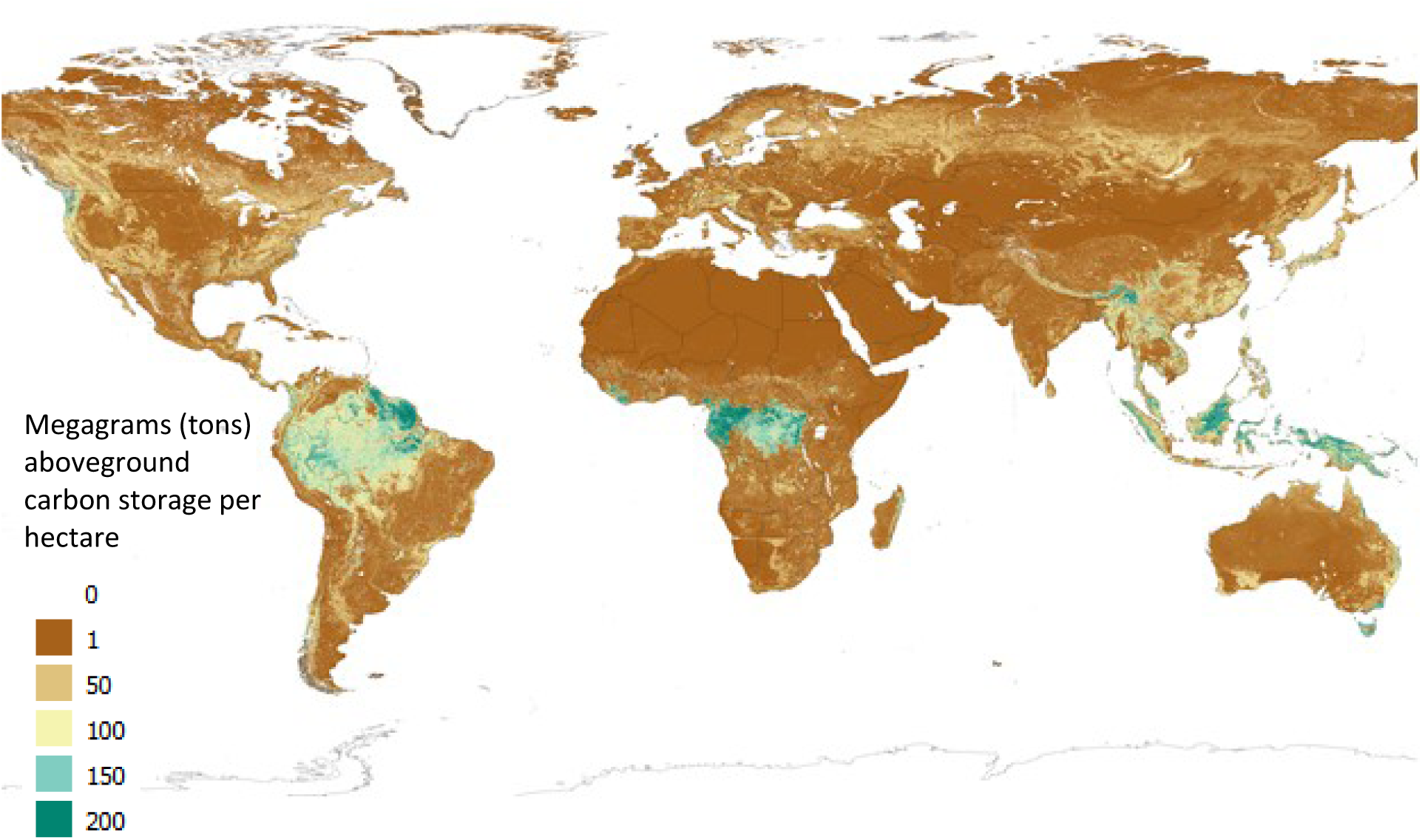
Aboveground carbon storage results from decision tree.

Visual inspection shows significant differences among the datasets, as shown in figure 3, highlighting how the map in figure 2 combines the best characteristics of each dataset.

**Figure 3:**
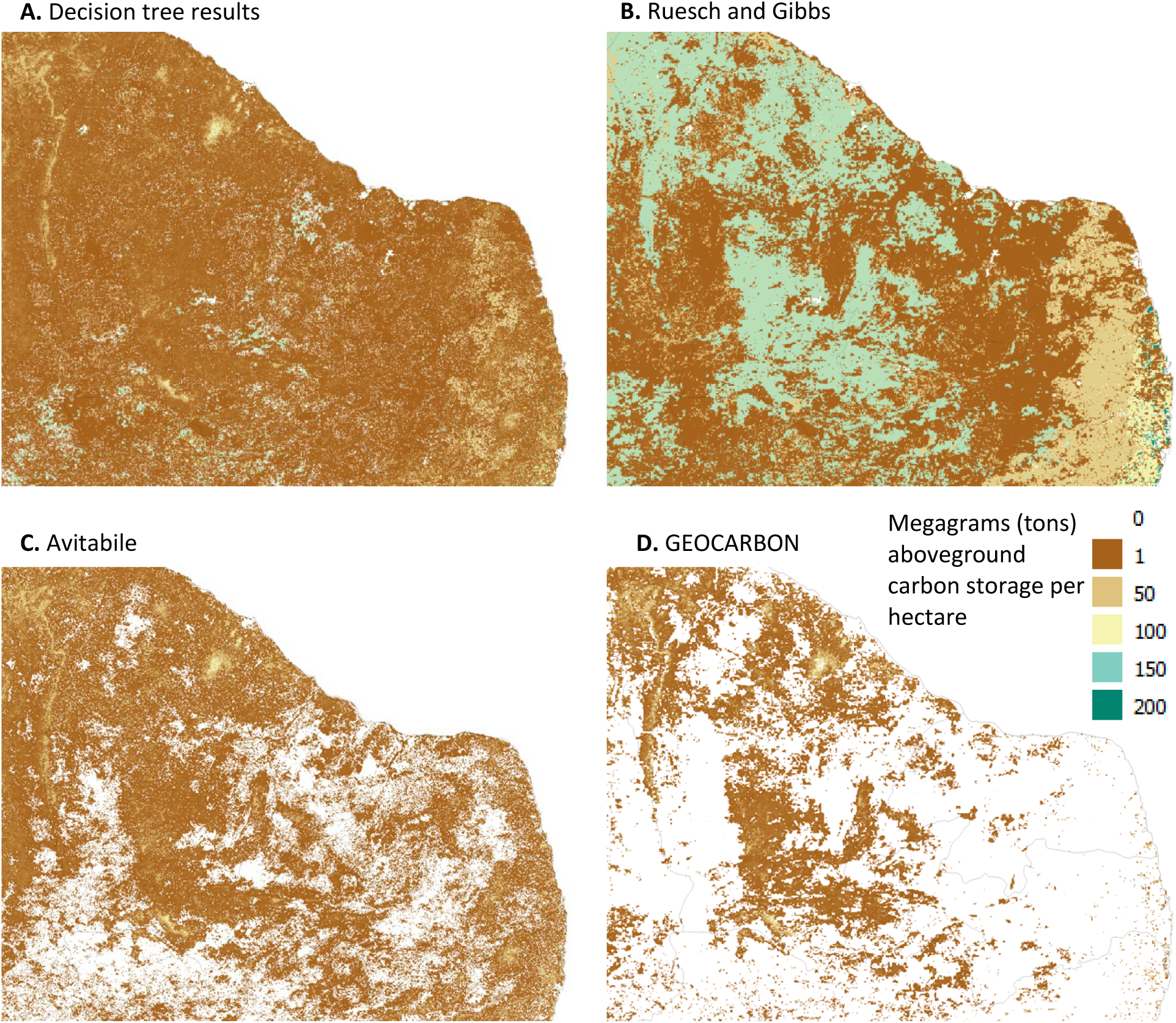
Aboveground carbon storage results compared to input for eastern Brazil.

Finally, of the two maps that are global in extent (decision tree results and Ruesch and Gibbs), summation of total carbon values shows significant differences. To calculate this, I first calculated the number of hectares present on each 30 arcsecond grid-cell (correcting for the spheroidal nature of the earth, following the method used by Doxsey-Whitfield et al. 2015). This map was then multiplied by the carbon per hectare results from above to generate total values. The summation of the decision tree result was 336.55 petagrams while the summation of Ruesch and Gibbs was 502.38 petagrams (49.27% higher than the results from the decision tree).

## Discussion

This work applied a simplified decision tree to multiple carbon storage datasets in order to obtain global coverage. The approach here could easily be improved by, for instance, applying a global application of the statistical method in Avitabile, though this has not yet been done publicly. Before such results are available, this product may be useful to conservation practitioners who use carbon storage as a metric of quality when prioritizing which areas to fund for conservation efforts.

## Data and code download

Results presented here are freely available and can be downloaded from http://justinandrewjohnson.com/data. The code used to generate these files is open-source and available at https://github.com/jandrewjohnson/harmonized_carbon_storage.

